# Building online genomics applications using BioPyramid

**DOI:** 10.1101/243378

**Authors:** Liam Stephenson, Yoshua Wakeham, Nick Seidenman, Jarny Choi

## Abstract

**BioPyramid** is a python package, which can serve as a scaffold for building an online genomics application. BioPyramid contains a number of components designed to reduce the time and effort in building such an application from scratch, including gene annotation, dataset models and visualisation tools. The user can rapidly deploy a data portal with the example dataset included, and start customising components as required. BioPyramid is implemented in python and javascript and freely available at http://github.com/jarny/biopyramid.

## Introduction

In the past decade, we have seen a huge increase in the number of high throughput genomics datasets produced, such as RNA-Seq datasets, primarily driven by cheaper and more accessible technologies. Following this trend has also been an increase in the number of applications which act as online portals to the datasets. Often designed for the bench biologists, these applications enable easy access and analyses of the datasets they host, enhancing their value and acting as hypothesis generators. Some examples include Haemosphere [7], Immgen [8], Stemformatics [10] and GeneExpression Commons [9]. These applications also contain a set of visualisation tools such as gene expression plots or heatmaps.

The ability to quickly create a custom application for data analysis and visualisation enhances collaboration and sharing of ‘omics’ projects. Typical example use-cases may be to host private datasets within a collaborative network, or to bring together related datasets from disparate sources so that many users can access them easily, or to try new analysis or visualisation tools within groups. However building such applications from scratch can require considerable time and effort.

We have developed **BioPyramid** as a python and javascript based framework for developing an online genomics application. It can be used as a starting scaffold or a template, to quickly get started with building a data portal. While many online genomics data portals exist, we are not aware of any toolkits or frameworks designed to help building such an application within the bioinformatics context.

**Figure 1.**
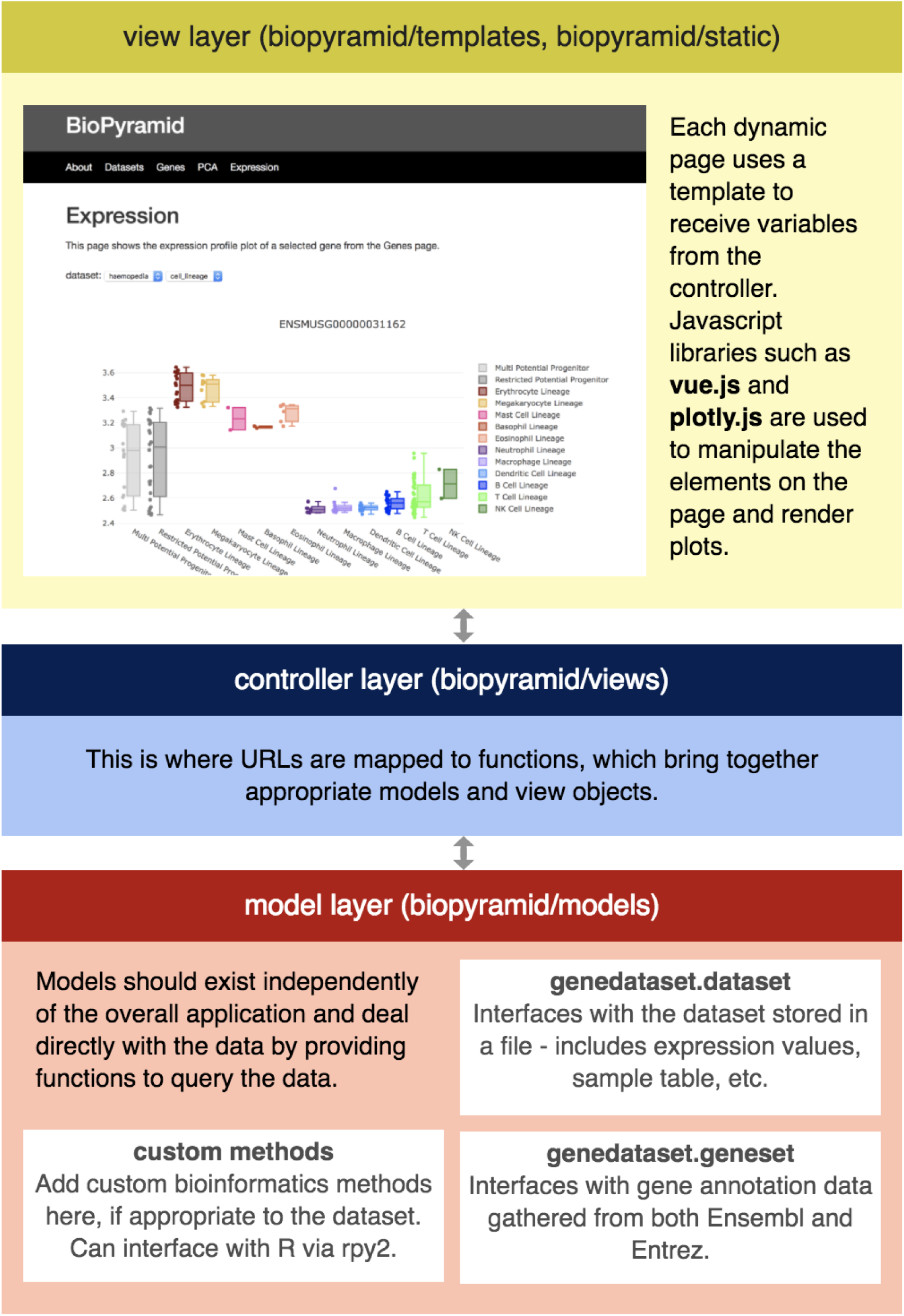
The design of BioPyramid and its components. This figure also shows what the default expression profile page looks like.

## Components and Customisation

BioPyramid is built upon the python pyramid framework [3], which provides a flexible and scalable framework for a web application. It uses a python package called genedataset [6], which was custom developed to to handle both dataset and geneset models. Within this package, Dataset is a model for storage and handling of expression values, sample table and other relevant data, while Geneset is a model that stores gene annotation data collected from both Ensembl [11] and NCBI Entrez [5].

Both models take advantage of the read-only nature of the application, and use the HDF file format [1] to store all the different data types in one file. This greatly improves data retrieval speed, compared to reading text files of expression matrices for example. For visualisation, BioPyramid comes with example pages showing fully interactive scatter and bar plots, using the established javascript library plotly [2]. It also comes with an example dataset [7], which comprises of gene expression data across a comprehensive range of murine haeamatopoetic cell types.

The developer can choose to include or exclude any of these components quite easily. The code comes with ample documentation and comments to help developers customise the application. The philosophy behind the design was to create a minimal working product with clear instructions on how different pieces fit together, rather than provide lots of ready-made custom options, in order to provide maximum flexibility. Hence even though these components are geared towards supporting gene expression datasets, the framework itself provides a useful basis for many other data types and workflows.

## Implementation and Availability

BioPyramid is written in python and javascript. The only pre-requisite to install python. BioPyramid is freely available with MIT license [4] from github.com/jarny/biopyramid.

## Acknowledgments

Thanks to Rowland Mosbergen and Christine Wells for helpful discussions, and to Steve Englart for testing. This work is in part supported by a Science and Industry Endowment Fund (SIEF) grant, funding from CSL and from Stem Cells Australia.

